# Stiffness of vascular smooth muscle cells from monkeys studied using atomic force microscopy

**DOI:** 10.1101/2020.12.12.422531

**Authors:** Yi Zhu

## Abstract

Vascular smooth muscle cells (VSMC) are the main cellular components of blood vessel walls and bear external mechanical forces caused by blood flow and pressure. In this report, we have verified the following hypothesis through experiments: The increase in VSMC stiffness may be mainly due to changes in vascular stiffness due to aging. Although aging enhances the stiffness and adhesion of VSMC, there is no significant difference in apparent elastic modulus and adhesion between the VSMC obtained by male and female. The effect of aging through the ECM-integrin-cytoskeleton axis is related to increased VSMC stiffness and matrix adhesion rather than gender.

## 1. Introduction

Most prior studies have utilized either indirect measurements of vascular stiffness or examined isolated vessel preparations. Additionally, most prior work has focused on the extracellular matrix (ECM) or endothelial control, as the source of the changes in vascular stiffness (Lakatta E.G., et al., 2002a and 2002b; Mitchell G.F., et al., 2004 and 2007; Haddock R.E., et al., 2005; Qiu et al., 2007). These ECM changes were primarily seen as an increase in total collagen and a decrease in elastin (Qiu H. Y., et al., 2007; Hong Z.K., et al., 2015). VSMCs are the main cellular components of blood vessel walls and bear external mechanical forces caused by blood flow and pressure. The aging process reduces the number of VSMCs in arteries, and the remaining VSMCs are enlarged in cell size to work harder (Briones A. M., et al., 2010; Lee H.Y., et al., 2010). The single VSMC mechanical characteristics and behavior is chiefly considered to play a key role in the development of vascular diseases (Zhu Y., et al., 2018 and 2019). Cytoskeletal action and role in VSMCs include contractile filament cycling for the actin and myosin cross-bridge and remodeling of cytoskeletal architecture and construction to employ and perform the mechanical forces (Levental K.R., et al., 2009; Niland S., et al., 2011). The cytoskeletal behaviors flexibly and livelily support and sustain VSMC elastic and adhesive properties to respond and regulate vascular physiological functions. More importantly, the cytoskeleton is physically and functionally participated and joined to the cell adhesive process. VSMC imposes the regulation and control of ECM assembly, composition and architecture through cell adhesion, and they are likely playing an important role in the ECM stiffening associated with aging. Meanwhile, VSMCs also can conversely alter their signaling state to reflect the rigidity of the ECM through adhesions to the ECM cells. Thus, the relationship is made complex by the changing and lively two-way interaction between ECM-adhesion site-cytoskeleton (Staiculescu M.C., et al., 2014). The AFM probe deflection and applied force can be used to determine cellular elasticity by an AFM probe indenting the surface of the cell (Sader J. E., et al., 1999; Labernadie A., et al., 2010). The AFM probe is functionalized with a ligand, and then the adhesion of the AFM probe to the cell surface can be used to evaluate cell adhesive interactions and this has been a successful application for studies of VSMC-ECM interaction (Sun Z., et al., 2005, 2008 and 2009; Hong Z., et al., 2015). For these studies we used atomic force microscopy and imaging to estimate and evaluate Young’s modulus of elasticity and adhesion in aortic smooth muscle cells derived from groups of old and young primates. Our studies have revealed some very novel findings regarding changes in smooth muscle cell mechanical characteristics that occur during aging. We had observed that smooth muscle cell stiffness and adhesion to the extracellular matrix is significantly increased in old monkey cells compared to those of young counterparts.

An important feature of this investigation was the use of the non-human primate model (*M. fascicularis*). The primate model has a closer phylogenetic proximity to humans and also exhibits an aging process that occurs more gradually over 20-30 years, vs 1-2 years in rodents. The mechanical properties of the VSMCs were measured directly in vitro using AFM. The apparent Young’s modulus was significantly increased in VSMCs from old male and female monkeys compared to young male and female counterparts. Moreover, the measured adhesion force was significantly increased in VSMCs from old male and female monkeys compared to young male and female monkeys. However, the apparent elastic modulus and adhesion force were not significantly different between male and female in the same aging group. The pan actin intensities of VSMCs using immunostaining analysis and the cell sizes of VSMCs from the old group were higher than those from the young group both in male and female monkeys. The apparent values were not significantly different between male and female in the same aging. The cell heights were not significantly changed due to aging and gender difference. Our results demonstrated that the intrinsic mechanical properties of VSMCs contribute to the increased vascular stiffness associated with aging and without gender.

## 2. Materials and Methods

### 2.1 Animal model

In this study young (6.4±0.1 years) and old (25±0.4 years) monkeys (Macaca fascicularis) (n=3/group) were used and maintained in the Guide for the Care and Use of Laboratory Animals (NIH 85-23, revised 2011).

### 2.2 VSMC isolation and culture

VSMCs from thoracic aorta of monkeys were enzymatically isolated and used at passages 2 to 4. All external fat, connective tissue, adventitia and endothelial layer were carefully detached and removed. The prior publications provided the detail methods (Qiu H.Y., et al., 2010; Zhu Y., et al., 2012; Sehgel N.L., et al., 2013).

### 2.3 AFM imaging, VSMC stiffness and force measurement

The topographical images of VSMC in this chapter were operated in contact mode by an AFM instrument, which is a Bioscope System (model IVa, Veeco Mertrology Inc.) mounted on an Olympus IX81 microscope (Olympus Inc.), and the AFM probe scanned across the cell surface (35 to 40 mm/s) with a tracking force of 300~500 pN. Transverse VSMC stiffness was measured in force mode using silicon nitride cantilevers (Trache A., et al., 2008; Hong Z.K., et al., 2012; Sehgel N.L., et al., 2015), and the AFM probe was repeated to approach and retract from the cell surface at 0.5 Hz (tip speed 800 nm/s) to collect 15-20 force curves from the same site on each VSMC (30-40 s of recording). 10-20 cells were randomly selected and the AFM probe indented at a site midway between the nucleus and cell margin (150-400 force curves per cell) for each experiment. Young’s modulus theory was applied to interpret and translate the recorded approach-rapture force curve into the AFM force-indentation description and statement for the quantification of VSMC elasticity. The calculation of the elastic modulus was:

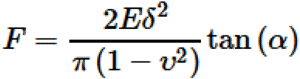

where the indentation force (F) was stated and described using Hooke’s law (F =κΔx, κ and Δx denote the AFM probe’s spring constant and the probe’s apparent deflection). The indentation depth (δ) is identified from the difference in the AFM piezo movement in z direction and the AFM probe deflection. E is the Young’s modulus of experimental cell as the value of elasticity, and ν denotes 0.5 for cell as the Poisson ratio. The numerical α is the semi-included angle of the cone for a pyramidal tipped probe and determined by the probe shape (MLCT, Veeco Mertrology Inc., Santa Barbara, CA) (Sun Z, et al., 2012; Hong Z., et al., 2014). An authoritative software NforceR is often applied to quantitatively analyze the apparent force-distance curves (Trzeciakowski J.P. et al., 2006).

β1 integrin was identified on the VSMC surface using AFM probes labeled with fibronectin (1mg/ml) (Invitrogen, Grand Island, NY). Probes were operated in force-mode and repeated to indent and retract from the cell surface and specific adhesions detected as sharp shifts in the retraction curves using automated software (NforceR) (Trzeciakowski, J.P., et al., 2006). With each probe, 10 randomly selected cells were sampled 50-60 times to collect approximately 500-600 force curves for analysis.

### 2.4 Immunostaining analysis

Isolated VSMCs were cultured and grown for 48 hours on glass substrate dishes under 37^°^C and 5% CO_2_ environment, and then fixed in 4% formaldehyde. The fixed cells were permeabilized with 0.1% Triton-X 100 for 5 minutes, rinsed and incubated with the primary antibodies overnight. Rinsed and washed 6 times using an antibody buffer, the cells were stained with the fluorescent secondary antibody. The cells were imaged by confocal microscopy for pan actin and α5β1-integrin.

### 2.5 VSMC cell area and height measurement

Analyzed and measured VSMC cell area and height from AFM deflection images using the bioscope software (a Bioscope System, model IVa, Veeco Mertrology Inc.).

### 2.6 Statistical analysis

Data are described as mean ± SEM for the number of samples reported in each figure legend. Statistically significant differences between young and old monkeys were identified using Student’s t-test. A value of P<0.05 was considered significant.

## 3. Results

### 3.1 VSMC AFM and confocal microscopic image and topography

To study VSMC physical character, we first observed and compared single cell area and height. From AFM image data (Figure 1), the male monkey cell area of old (10 cells from 3 animals) vs. young (10 cells from 3 animals) was 10,317 ± 24 μm^2^ vs. 10,167 ± 9 μm^2^ and the male monkey cell height of old (17 cells from 3 animals) vs. young (10 cells from 3 animals) was 3366 ± 214 nm vs. 3518 ± 124 nm. And the female monkey cell area of old (10 cells from 3 animals) vs. young (10 cells from 3 animals) was 10,285 ± 30 μm^2^ vs. 10,183 ± 8 μm^2^ and the female monkey cell height of old (14 cells from 3 animals) vs. young (14 cells from 3 animals) was 3171 ± 134 nm vs. 3366 ± 175 nm, respectively. The cell area of old monkeys was larger than that of young monkeys (P<0.05), and there was no significant difference between male and female in both old and young. Despite the differences of monkey age and gender, the cell height did not show a significant difference. The VSMCs of thoracic aorta from subject monkeys were also investigated by confocal microscopy, pan actin data was listed in Figure 2. From the immunostaining intensity analysis and comparison, the old male monkey VSMCs (8 cells from 3 animals) were higher than those of young male monkey (12 cells from 3 animals), and the old female monkey VSMCs (12 cells from 3 animals) were higher than those of young female monkey (22 cells from 3 animals), but in the same age group, male and female cells were not different in fluorescent intensity. Moreover, immunostaining intensity for α5β1-integrin in cultured VSMCs from thoracic aorta of monkeys was shown in Figure 3. The old male monkey VSMCs (11 cells from 3 animals) were higher than those of young male monkeys (17 cells from 3 animals), and the old female monkey VSMCs (19 cells from 3 animals) were higher than those of young female monkeys (16 cells from 3 animals), respectively. Interestingly, in the same age group, male and female cells were not different in fluorescent intensity.

**Figure. 1.**
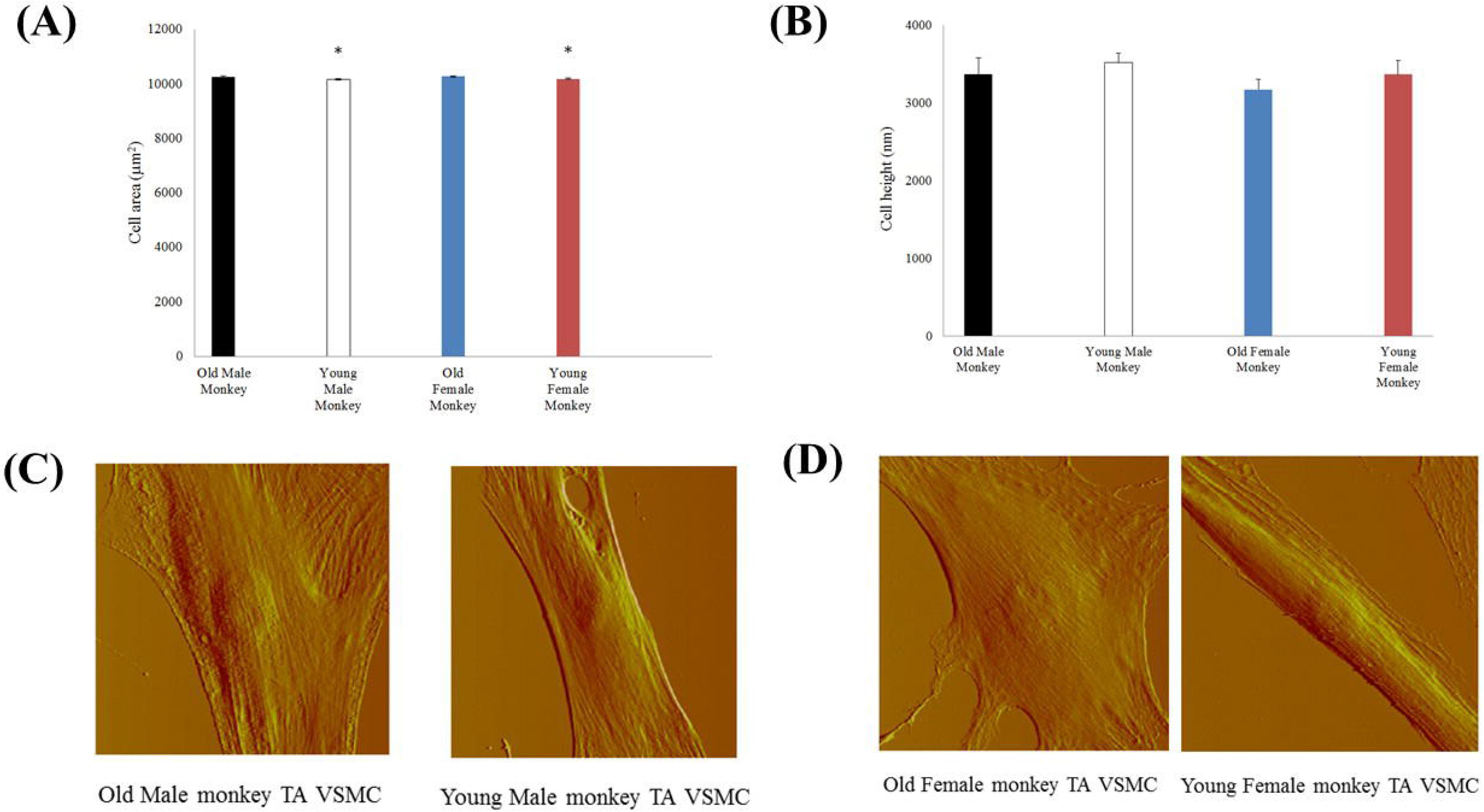
Topographic characterizations of old and monkey VSMCs. (A) VSMC area from thoracic aorta of old (male n=15, female n=11) and young (male n=10, female n=10) monkeys. *, P< 0.05 old versus young monkeys. p>0.05 male vs female. (B) VSMC height from thoracic aorta of old (male n=17, female n=14) and young (male n=10, female n=14) monkeys. P> 0.05 old versus young monkeys. p>0.05 male vs female. (C) AFM deflection images of thoracic aorta vascular smooth muscle cells from old and young male monkeys. (D) AFM deflection images of thoracic aorta vascular smooth muscle cells from old and young female monkeys.

**Figure. 2.**
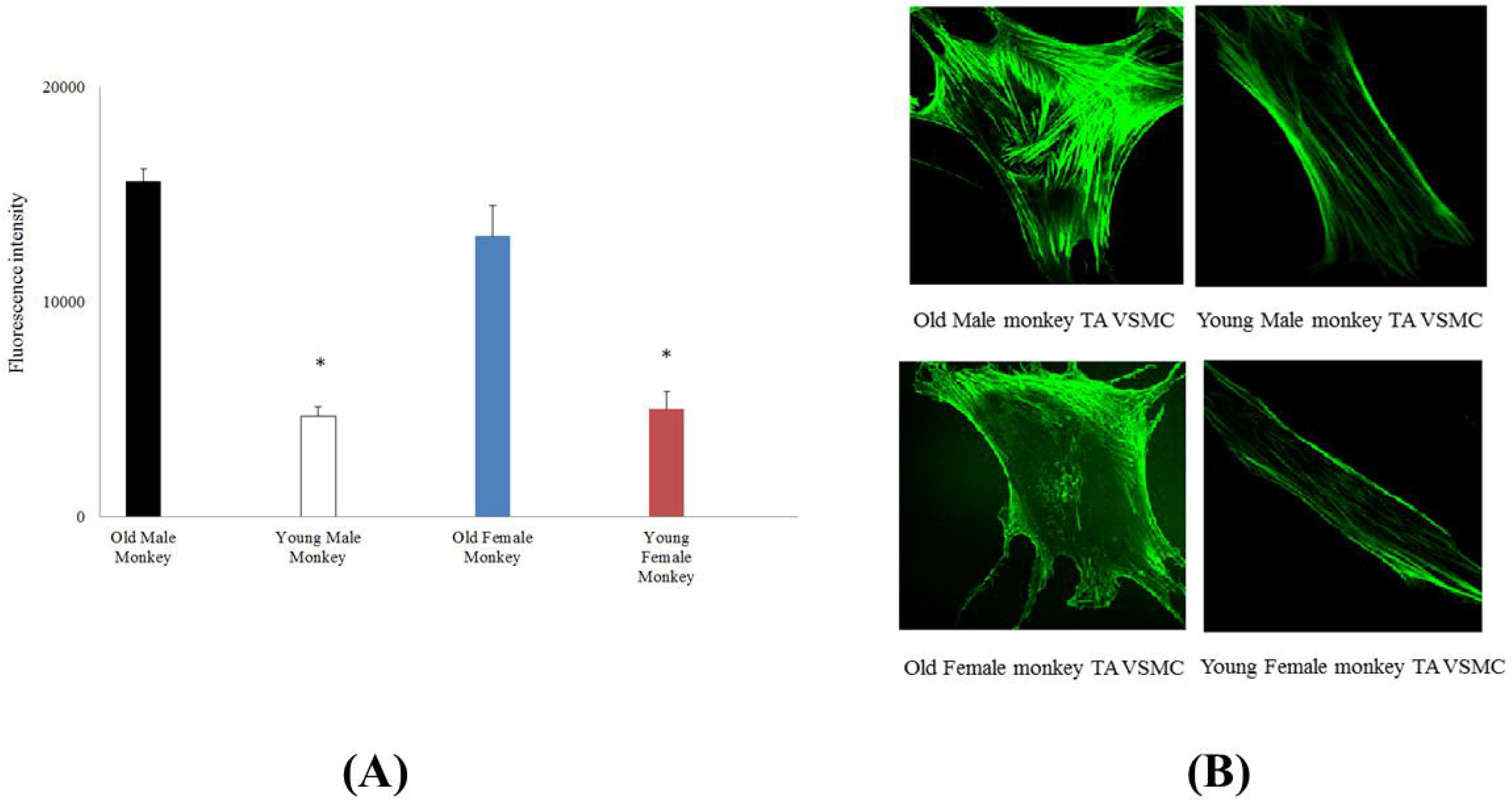
(A) Immunostaining intensity for pan actin in cultured VSMCs from thoracic aorta of old (male n=8, female n=12) and young (male n=12, female n=22) monkeys. *, P< 0.05 old versus young monkeys. p>0.05 male vs female. (B) Confocal microscopic images of pan actin for thoracic aorta VSMCs from old and young of male and female monkeys.

**Figure. 3.**
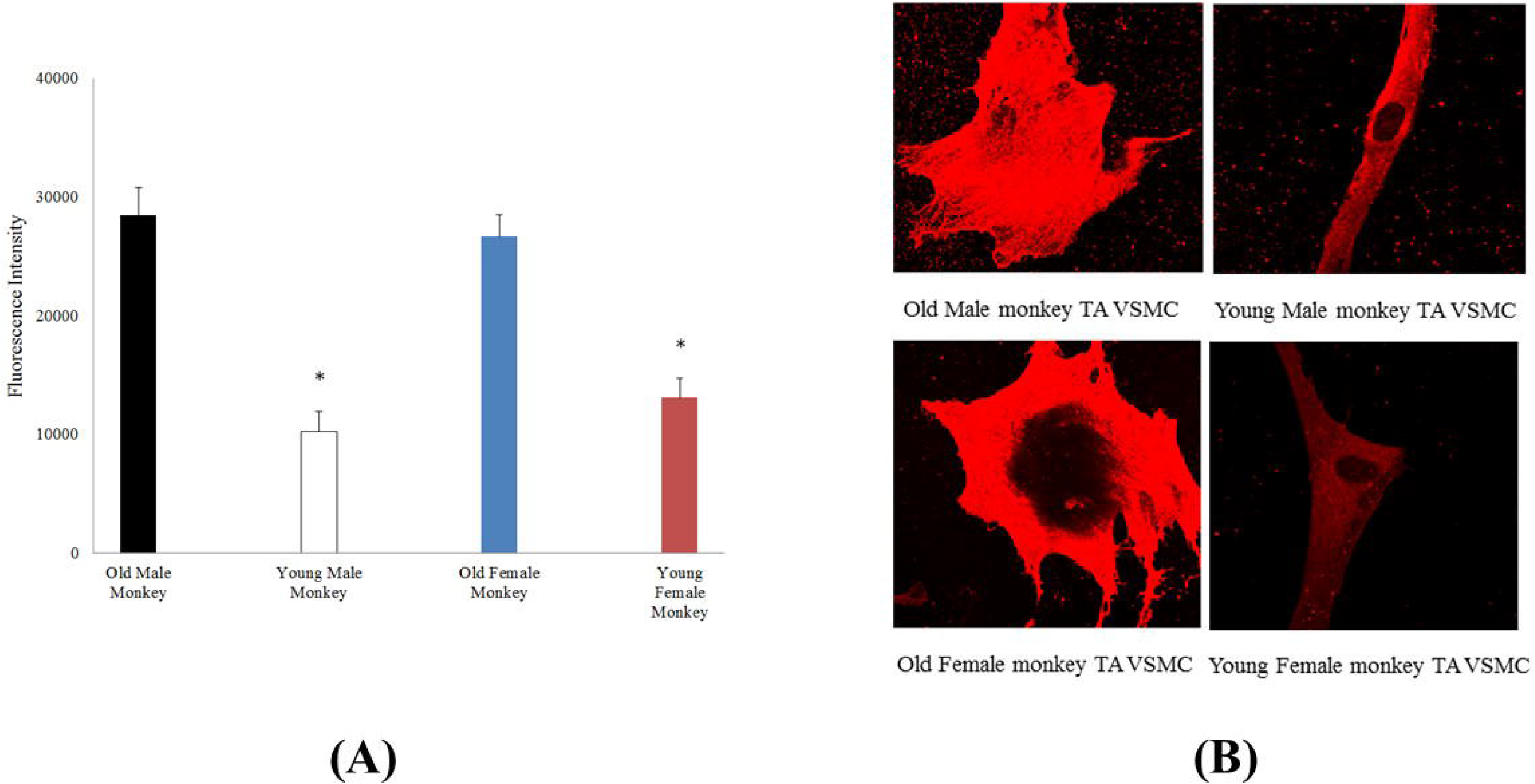
(A) Immunostaining intensity for α5β1-integrin in cultured VSMCs from thoracic aorta of old (male n=11, female n=19) and young (male n=17, female n=16) monkeys. *, P< 0.05 old versus young monkeys. p>0.05 male vs female. (B) Confocal microscopic images of α5β1-integrin for thoracic aorta VSMCs from old and young of male and female monkeys.

### 3.2 Elasticity and ECM protein-integrin adhesion

Single VSMC elasticity was measured using AFM nanoindentation. As shown in Figure 4, the apparent elastic modulus was significantly higher in VSMCs from old (42.6±3.4 kPa; 11 cells from 3 animals) compared with young (11.3±1.6 kPa; 6 cells from 3 animals) male monkeys. Moreover, the apparent elastic modulus was significantly higher in VSMCs from old (50.6±4.6 kPa; 7 cells from 3 animals) compared with young (12.8±1.3 kPa; 8 cells from 3 animals) female monkeys. These elasticity data demonstrate that VSMCs from old monkeys are inherently stiffer than those from young monkeys. In addition, AFM probes coated with fibronectin to measure the adhesion behavior of old versus young VSMC integrin α5β1 (Figure 4), and adhesive forces were also enhanced in old VSMCs, as evidenced by their increased adhesion to FN-coupled AFM probes. The measured force was significantly increased (p<0.05) in VSMCs from old male (77.6±12.4 pN; 11 cells from 3 animals), and old female monkeys (89.5±7.8 pN; 11 cells from 3 animals) compared to young male (43.5±3.8 pN; 6 cells from 3 animals), and young female monkeys (54.2±6.0 pN; 6 cells from 3 animals). In the same age group, male and female cells were not also different in stiffness and adhesive force.

**Figure. 4.**
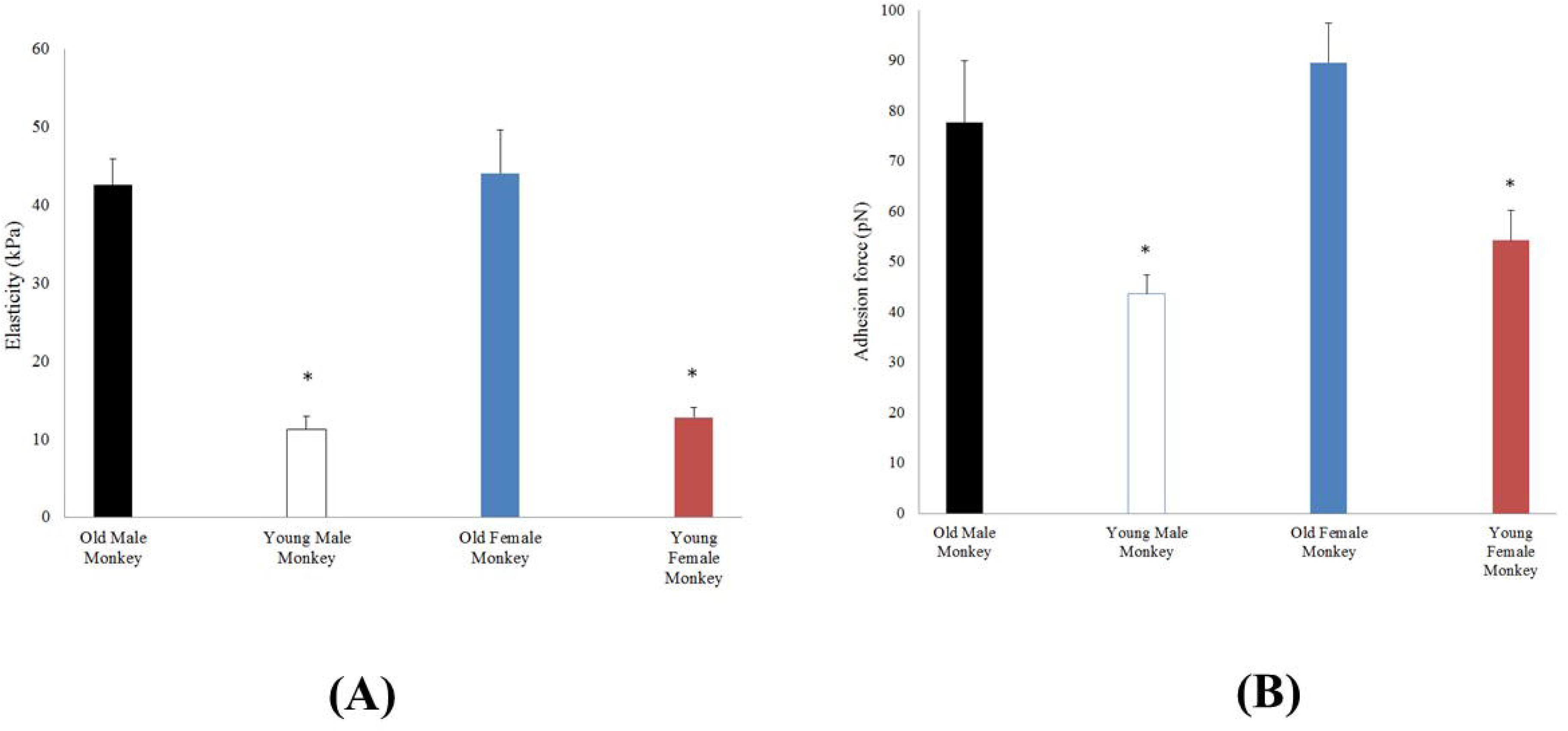
(A) Elasticity values measured by AFM in VSMCs from thoracic aorta of old (male n=11, female n=10) and young (male n=6, female n=8) monkeys. *, P< 0.05 old versus young monkeys. p>0.05 male vs female. (B) α5β1-integrin adhesion to the extracellular matrix (ECM) protein fibronectin (FN) on the surface VSMC from thoracic aorta of old (male n=11, female n=11) and young (male n=6, female n=6) monkeys by AFM adhesion force measurement. *, P< 0.05 old versus young monkeys. p>0.05 male vs female.

## 4. Discussion

Our prior research showed the abundance of alpha-smooth muscle actin in cytoskeleton of the old monkey VSMCs (Qiu H.Y. et al., 2010; Zhu Y. et al., 2012). From the AFM image data, the area of single VSMC is significantly higher in old compared to young monkeys, thus the physical character of single old monkey VSMC is different from that of young one. A single vascular smooth muscle cell from male monkeys (Macaca fascicularis) was measured with AFM indentation method, and the old VSMC is obviously harder than the young one in elasticity. Meanwhile, the alpha-SMA of cytoskeleton mainly contributes to the inherent mechanical feature of VSMC (Figure2B).

The adhesion force significantly enhanced in old compared to young monkeys due to old monkeys’ higher expression of β1-integrin (Qiu H.Y. et al., 2010; Zhu Y. et al., 2012). Fibronectin specifically binds with β1-intergrin (Parsons J.T., et al., 2010; Staiculescu M.C., et al., 2013). The α-SMA is an important component of the ECM-integrin-cytoskeltal axis responsible for mechanosensation, -transduction and -transmission (Geiger B., et al., 2001; Galbraith C.G., et al., 2002 and 2007). Meanwhile, the β1-integrin, which is one of transmembrane proteins for the interaction between the ECM protein and the cytoskeleton, provides an important mechanical role and association with the extracellular environment (Schwartz M.A., et al., 2002; Staiculescu M.C., et al., 2014; Han S.J. et al., 2015). Interestingly, the apparent values of stiffness of VSMCs between male and female of young monkeys were closed. In summary, we demonstrate that aging is associated with increased VSMC stiffness and cell-matrix adhesion via the ECM-integrin-cytoskeltal axis responsibility rather than gender.

